# A novel class of tiny box C/D-like RNAs reveals hidden complexity in the mouse box C/D RNA repertoire

**DOI:** 10.64898/2026.07.28.740266

**Authors:** Caetano Tomas, Grand Baptiste, Hebras Jade, Feat Michelle, Zytnicki Matthias, Cavaillé Jérôme

## Abstract

In recent years, the catalog of mammalian box C/D small nucleolar RNAs (SNORDs) has expanded considerably, with several hundred members now annotated in humans. Despite being a well-characterized family of antisense small noncoding RNAs, the full diversity of mammalian box C/D SNORDs remains incompletely understood. To address this, we combined deep sequencing of Fibrillarin- and Snu13-associated RNAs with systematic mining of a large collection of publicly available small RNA-seq datasets spanning diverse mouse tissues and cell types. This integrative approach uncovered an unexpectedly large repertoire of box C/D-like RNAs, comprising more than 20 000 distinct species. These RNAs display the defining hallmarks of intronic SNORDs but are expressed at low levels and exhibit poor evolutionary conservation. Strikingly, approximately 70% are only 20-50 nucleotides long, making them substantially shorter than canonical SNORDs. We designate this previously unrecognized class of box C/D-like RNAs as tiny SNORDs (tiSNORDs). Functional characterization of a representative 36-nucleotide tiSNORD demonstrates that, despite its reduced size, it retains nucleolar 2′-O-ribose methylation guide activity. The discovery of this previously unrecognized class of box C/D-like RNAs substantially expands the known repertoire of SNORDs and raises fundamental questions regarding their biogenesis, functional potential and evolutionary significance.

## Introduction

Box C/D small RNAs are part of ribonucleoprotein (RNP) complexes composed of three scaffold proteins (Snu13, Nop56 and Nop58) and a 2’-O-methyltransferase (Fibrillarin). Identified in both eukaryotes and archaea, they represent one of the best-known families of antisense small noncoding RNAs (1–5), with proposed ancestral relationships to the bacterial TIGR system and IS110 RNA-guided transposases (6, 7).

As their name suggests, all members contain two conserved short sequence motifs, the C (RUGAUGA) and D (CUGA) boxes, typically located near the 5′ and 3′ termini, respectively, as well as internal variants known as the C′ and D′ boxes. Apart from U3/SNORD3, U8/SNORD118 and U13/SNORD13, which are transcribed independently from their own promoters, all mammalian box C/D small RNAs are post-transcriptionally processed through exonucleolytic trimming of Pol-II transcribed host-introns and, depending upon accumulation of RNPs in the nucleoli or Cajal bodies, they are classified as small nucleolar RNAs (SNORD) or small Cajal body-specific RNAs (scaRNAs), respectively (8).

One key structural feature of box C/D RNPs is the presence of a kink-turn (k-turn) motif, formed by the conserved arrangement of the C- and D-box elements. The k-turn consists of a three-nucleotide bulge flanked by a short stem on one side and a non-canonical stem containing two conserved sheared G base pairs on the other side. This architectural RNA element introduces a kink in the helical axis and is directly recognized by SNU13 (also known as 15.5 kDa, NHPX or NHP2L1), which nucleates the stepwise assembly of the RNP and contributes to its stability, subcellular localization and functional activity (3, 9–13).

The D and D′ boxes also play a central role in antisense-mediated mechanisms. Indeed, the vast majority of box C/D small RNAs typically harbor one, and occasionally two, phylogenetically conserved antisense sequence(s), positioned immediately upstream of the D or D′ boxes. By tethering the RNP to their complementary RNA sequences through perfect base-pairings, these antisense elements, typically 10-20 nt in size, guide Fibrillarin to transfer a methyl group onto the ribose of the nucleotide paired to the fifth nucleotide upstream of the D or D’ box. SNORDs direct site-specific 2′-O-methylation of RNA Polymerase I-transcribed ribosomal RNAs (rRNAs) and RNA Polymerase III-transcribed U6 spliceosomal snRNA, whereas scaRNAs guide modifications of RNA Polymerase II-transcribed U1, U2, U4, and U5 spliceosomal snRNAs (14–16). Of note, RNA Polymerase III-transcribed tRNAMet can be targeted for ribose methylation by either a SNORD or a scaRNA (17). Beyond this main role in specifying 2’-O-methylations, a few SNORDs also ensure essential “chaperon-like roles” in the nucleolytic processing of rRNA precursors and pre-ribosome assembly (18–23) as well as in the synthesis of two N4-acetylcytidines in 18S rRNA (24–27). Finally, emerging evidence suggests that some SNORDs may contribute to diverse aspects of gene expression, particularly in the post-transcriptional regulation of pre-mRNAs, although their precise modes of action remain largely unknown (reviewed in (28–31).

Over the past 50 years, the repertoire of human SNORD has grown steadily. In the 70’s-80’s, the sequences of the most abundant ones - U3/SNORD3 (32, 33), U8/SNORD118 (34, 35) and U13/SNORD13 (35) - were directly determined by *in vivo* [^32^P] labeling followed by enzymatic RNA fingerprints, or direct RNA sequencing. It was however in the 1990s that the complexity of the SNORD family was fully appreciated thanks to the sequencing of cDNA libraries made from nucleolar-enriched RNAs (16), or by systematic searching for conserved intronic sequences in protein-coding genes, notably those linked to ribosome synthesis and function (36–43). Sequencing of cDNA libraries prepared from size-fractionated RNAs (“RNOmics”) or RNAs recovered from immunoprecipitation of Fibrillarin also identified additional members, notably many tandemly-repeated, imprinted SNORD genes showing predominant expression in the brain (44–46). More recently, comparative genomic analyses (47, 48) and next-generation sequencing (NGS)-based methods (49–52) have completed the inventory of SNORDs in the human genome, which is now believed to encode a few hundred box C/D RNAs (n∼500), including canonical SNORDs and their paralogs, as well as pseudogenes and poorly characterized box C/D-like RNA species whose functions, if any, remain to be determined (53, 54).

Here, we used deep sequencing of Fibrillarin-associated RNAs from mouse tissues, together with systematic mining of publicly available small RNA-seq datasets, to show that the repertoire of box C/D RNAs is far more complex than previously appreciated. Specifically, we identify more than 20 000 novel box C/D-like RNA species that are lowly expressed and poorly conserved, with a substantial fraction (∼70%) measuring 20-50 nucleotides in length. We discuss the evolutionary and functional significance of this unexpectedly large class of novel box C/D RNA-like molecules, which we term tiny SNORD (tiSNORD).

## Results

In an effort to better characterize the complexity of the SNORD repertoire in tissues, an area that has been relatively understudied, and to expand our understanding of their regulatory landscape, we performed immunoprecipitation (IP) experiments using an anti-Fibrillarin antibody (monoclonal 72B9) on whole extracts prepared from adult mouse brain, developing (E15.5) brain, placenta and whole body. The RNAs extracted from pellets were used to create Illumina libraries, which were sequenced to obtain approximately 83-129 million reads per library (Figure 1A). The IP procedure recovered the vast majority of known murine SNORDs, which accounted for 67-93% of annotated sequencing reads in different tissues (Figure 1B; Supplementary Table 1), thus demonstrating the robustness and specificity of our approaches. It should be acknowledged, however, that the observed abundance of SNORDs, as inferred from read counts, varied substantially among SNORD families. In particular, SNORD50 accounted for the majority of retrieved SNORD reads in all four tissues, a pattern that most likely reflects a bias arising from library preparation rather than true biological abundance.

**Figure 1.**
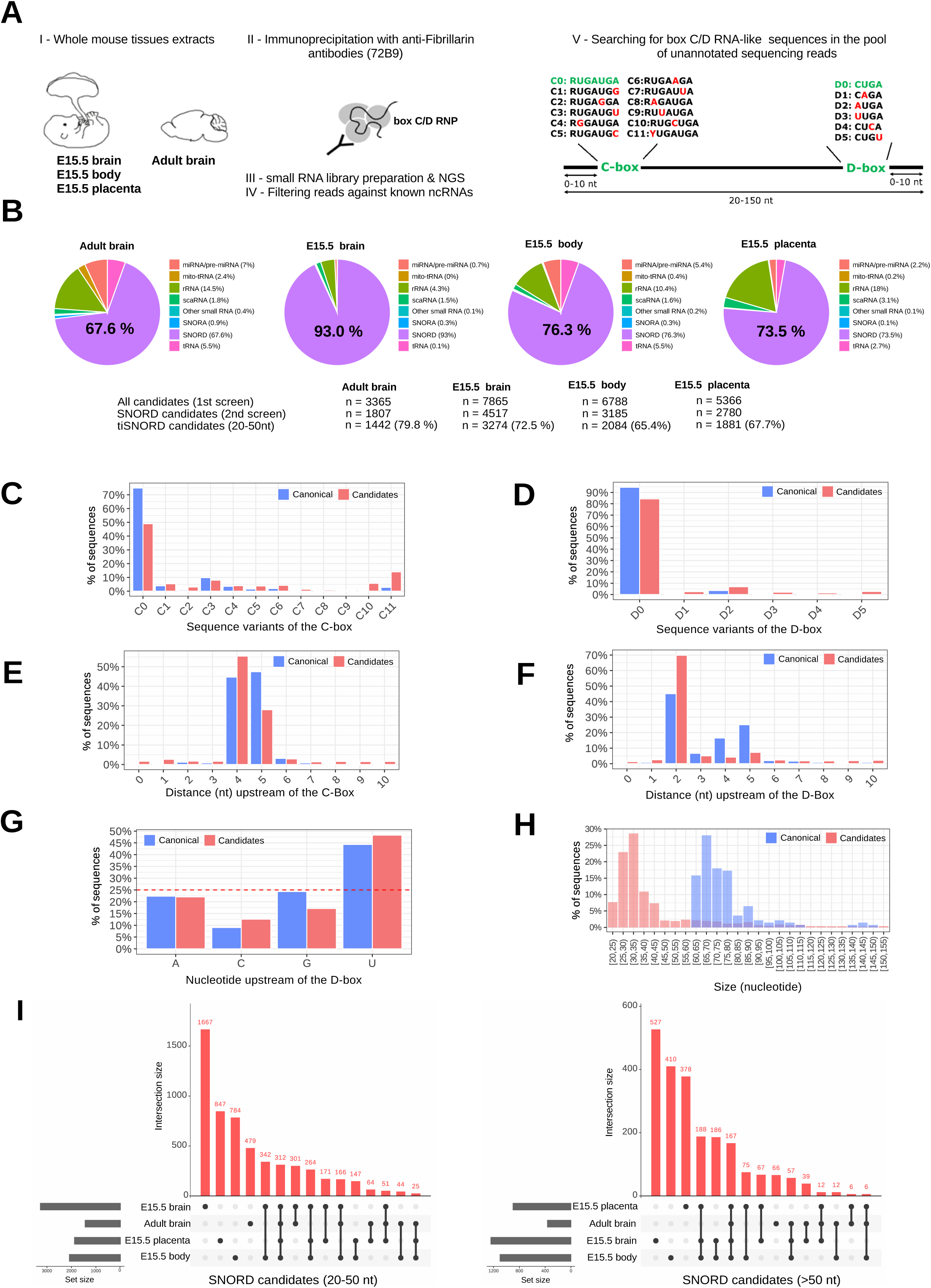
Identification of novel box C/D RNA-like RNAs in mouse tissues. (A) Following anti-Fibrillarin immunoprecipitation performed on whole-cell extracts from adult brain and from developing (E15.5) brain, trunk and placenta, Illumina sequencing reads (20-150 nt) generated from the pellet fraction that did not match any annotated ncRNAs were screened for the presence of C- and D-box motifs located no more than 10 nt from the 5′ and 3′ ends, respectively. The sequences of a few C- and D-box variants retained in our analyses are shown. (B) Top: Pie charts showing the relative proportions (%) of known ncRNA classes identified across the four mouse tissues analyzed. Bottom: The numbers of uncovered SNORD candidates (first and second screens) are shown, including the percentage of candidates ranging from 20 to 50 nt in size (tiSNORDs). (C) Relative proportions of C-box sequence variants (%) in unannotated sequencing reads (red) and canonical SNORDs (blue). (D) Relative proportions of D-box sequence variants (%) in unannotated sequencing reads (red) and canonical SNORDs (blue). (E) Relative proportion (%) of the positions of the C-box relative to the 5′ end of unannotated sequencing reads (red) and canonical SNORDs (blue). (F) Relative proportion (%) of the positions of the D-box relative to the 3′ end of unannotated sequencing reads (red) and canonical SNORDs (blue). (G) Overrepresentation of uridine residues upstream of the D-box in selected SNORD candidates (red) and canonical SNORDs (blue). The dashed red line indicates 25%, the proportion expected by random sampling. (H) Size (nt) distribution of selected SNORD candidate (red) and canonical SNORDs (blue). (I) UpSet plots showing the overlap of SNORD candidates between the four analyzed tissues, stratified by length (20-50 nt, left and >50 nt, right). (C-G): Adult mouse brain was selected as a representative example, while molecular profiles for the other three tissues are presented in Supplementary Figure 1.

Using sequencing reads that did not match any annotated RNAs, we then searched for RNAs between 20 and 150 nucleotides in length that were supported by at least two identical reads and, more importantly, carried C- and D-boxes positioned within 10 nucleotides of their 5′ and 3′ ends, respectively. In addition to the most commonly found C (C0; RUGAUGA) and D (D0; CUGA) sequence motifs, this analysis included several C- and D-box variants (C1-C8, C11; D1-D3), as well as mutated forms predicted to disrupt k-turn formation (C9-C10; D4-D5) and used as negative controls. Using this deliberately straightforward, low-stringency pipeline (Figure 1A), we retrieved 3365, 7865, 6788 and 5366 sequencing reads from pellets in adult mouse brain, developing (E15.5) brain, whole body and placenta, respectively (Figure 1B; Supplementary Table 2).

The large number of identified hits, collectively referred to hereafter as SNORD candidates, raises the possibility that many of them may represent “junk RNA,” reflecting background signals derived from RNA degradation and/or spurious transcription events captured by deep sequencing analyses. However, independent and complementary lines of evidence argue against this trivial interpretation and instead demonstrate that sequences of SNORD candidates exhibit non-random, reproducible features. These observations are illustrated for adult brain in Figure 1 (panels C-G), and comparable molecular profiles were observed across three developing tissues analyzed (Supplementary Figure 1). First, a substantial proportion of SNORD candidates contain the most represented C0 and D0 motifs, with the frequency of other sequence variants comparable to that of canonical SNORDs used as internal references for direct comparison (Figures 1C-D). Notably, dead variants unable to form GA:GA base-pairs essential for k-turn formation (C9-C10 and D4-D5) were underrepresented. Under the background hypothesis, this strong bias toward functional C- and D-boxes is difficult to explain. Second, a large fraction of the SNORD candidates displays the C-box at 4-5 nucleotides from the 5′-end and the D-box 2-5 nucleotides from the 3′-end (Figures 1E-F), as typically found for canonical SNORDs (16). Not only do these stereotyped positional end signatures argue against random sampling, but they also suggest that these SNORD candidates entered the same nuclear machinery responsible for the exonucleolytic trimming of canonical SNORDs. Third, approximately 50% of the SNORD candidates carry a uridine immediately upstream of the D-box, compared to the 25% expected by chance. This overrepresentation, which was not accounted for in our initial pipeline, is also a characteristic of canonical SNORDs (Figure 1G). Overall, these three observations indicate that SNORD candidates deviate from the nucleotide composition expected under random sampling and, collectively, they provide strong and unbiased experimental support for the classification of SNORD candidates as putative members of the box C/D small RNA family.

To minimize false-positive signals, this first list of SNORD candidates was subsequently refined by retaining only sequencing reads containing the most representative variants of the C- and D-box (C0, C1, C3, C4 and D0), with correct positioning relative to the 5′ and 3′ termini. This filtering step (2^nd^ screen) yielded 1807, 4517, 3185 and 2780 candidates in adult mouse brain, developing (E15.5) brain, whole body and placenta, respectively (Figure 1B; Supplementary Table 2). Strikingly, their size distribution differs markedly from that of canonical SNORDs, with 65-80% of the identified SNORD candidates ranging from 20 to 50 nt in length (Figure 1H). Collectively, we identified 7860 sequencing reads exhibiting hallmark SNORD features, including 2196 longer than 50 nucleotides and 5664 ranging between 20 and 50 nucleotides (Supplementary Table 2). Remarkably, only a limited subset of these SNORD candidates was detected across all four mouse tissues analyzed, suggesting possibly a degree of tissue specificity (Figure 1I).

As Fibrillarin shows little to no sequence-specific RNA-binding activity (55), the most likely hypothesis is that SNORD candidates are also bound by Snu13 or by as-yet unidentified Snu13-like proteins that directly interact with k-turn (56). To test this, we applied the same pipeline to published high-throughput datasets of RNAs recovered from Snu13 immunoprecipitation in mouse Hepa 1.6 cell lines (52). As shown in Figure 2A, we identified 18443 (1^st^ screen) and 7035 (2^nd^ screen) sequencing reads displaying the same structural features as those observed for Fibrillarin-associated RNAs in tissues, namely the correct positioning of the C- and D-boxes relative to the sequencing read termini (Figures 2B-C), the enrichment of the most common C- and D-box sequence variants (Figures 2D-E) and the presence of a uridine residue upstream of the D-box in approximately 50% of candidates (Figure 2F). Again, the size distribution of Snu13-associated SNORD candidates was strongly skewed toward short species, with 82% (5,790/7,035) ranging from 20 to 50 nt in length (Figure 2G), of which 26.1% (1516/5790) had already been identified in the Fibrillarin pellet (Figure 2H, left). A comparable overlap was also observed for longer SNORD candidates, with 28.4% (354/1245) shared between IP datasets (Figure 2H, right). Given the different biological sources (*in vivo* tissues vs *in vitro* immortalized cell lines), this degree of overlap is notable. Considering both Fibrillarin and Snu13 IP datasets, we identified 13,025 SNORD candidates, the vast majority of which (76.3%; 9,938/13,025) were only 20-50 nt in length

**Figure 2.**
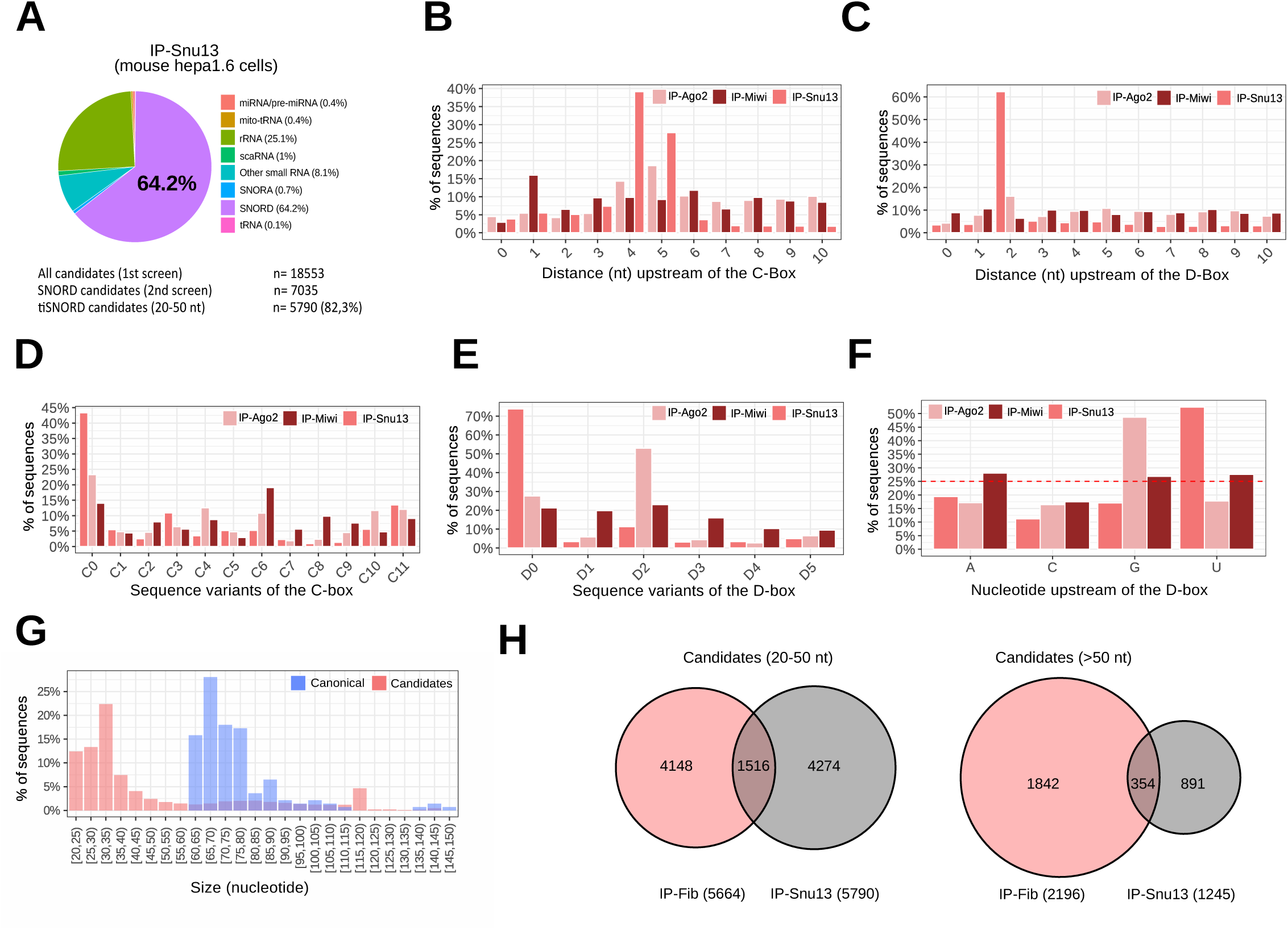
Identification of novel box C/D RNA-like RNAs in mouse Hepa 1.6 cells. (A) Top: Pie charts showing the relative proportions (%) of known ncRNA classes identified following anti-Snu13 immunoprecipitation in mouse Hepa 1.6 cells. Bottom: The numbers of uncovered SNORD candidates (first and second screens) are shown, including the percentage of candidates ranging from 20 to 50 nt in size (tiSNORDs). (B) Relative proportion (%) of the positions of the C-box relative to the 5′ end of unannotated sequencing reads following anti-Snu13 (mid pink), anti-Ago2 (pale pink) or anti-Miwi (dark pink) immunoprecipitations. (C) Relative proportion (%) of the positions of the D-box relative to the 3′ end of unannotated sequencing reads following anti-Snu13 (mid pink), anti-Ago2 (pale pink) or anti-Miwi (dark pink) immunoprecipitations. (D) Relative proportions of C-box sequence variants in unannotated sequencing reads following anti-Snu13 (mid pink), anti-Ago2 (pale pink) or anti-Miwi (dark pink) immunoprecipitation. (E) Relative proportions of D-box sequence variants in unannotated sequencing reads following anti-Snu13 (mid pink), anti-Ago2 (pale pink) or anti-Miwi (dark pink) immunoprecipitations. (F) Overrepresentation of uridine residues upstream of the D-box in selected Snu13-associated SNORD candidates. The dashed red line indicates 25%, the proportion expected by random sampling. (G) Size distribution (nt) of selected snu13-associated SNORD candidates (pink) compared with canonical SNORDs (blue). (H) Ven diagramms showing the overlap of SNORD candidates identified by anti-Fibrillarin and anti-Snu13 immunoprecipitation, stratified by RNA length (20-50 nt, left and >50 nt, right).

Such a large fraction of short RNA species raised the possibility that they may represent a subset of previously unrecognized small RNAs associated with components of RNA interference (RNAi) machinery. As reported in Figures 2B-F, this hypothesis can be ruled out, as no obvious enrichment of canonical SNORD signatures was observed following analysis of Ago2 immunoprecipitation in mouse embryonic fibroblasts (miRNA pathway) or Miwi immunoprecipitation in P4 testes (piRNA pathway). Altogether, our data support the existence of a previously underappreciated population of short box C/D-like RNAs, which we collectively term tiSNORDs (tiny SNORDs).

In order to expand the catalog of tiSNORDs, whose detection may have been masked by restricted tissue- or cell-type-specific expression profiles, we next applied the same bioinformatic pipeline to a large collection of publicly available small RNA-seq datasets derived from multiple and diverse adult and developing mouse tissues, as well as primary cell cultures or immortalized cell lines. The full list of examined datasets is provided in Supplementary Table 3. This strategy proved successful, enabling the identification of an additional set of more than 8000 members, bringing the total catalog to 21732 SNORD gene loci candidates, representing a total of 19342 distinct small RNA sequences (Figure 3A, Supplementary Table 4). Examples of SNORD signatures detected for embryonic stem cells, bone marrow and pancreatic αTC1 cells are given in Supplementary Figure 2. Among the tissues analyzed, tiSNORDs were especially abundant in the spleen and bone marrow, although the relevance of this enrichment remains unclear. Overall, these comprehensive analyses indicates that tiSNORDs are widespread and can be detected in standard small RNA-seq datasets without prior enrichment of box C/D RNP complexes.

**Figure 3.**
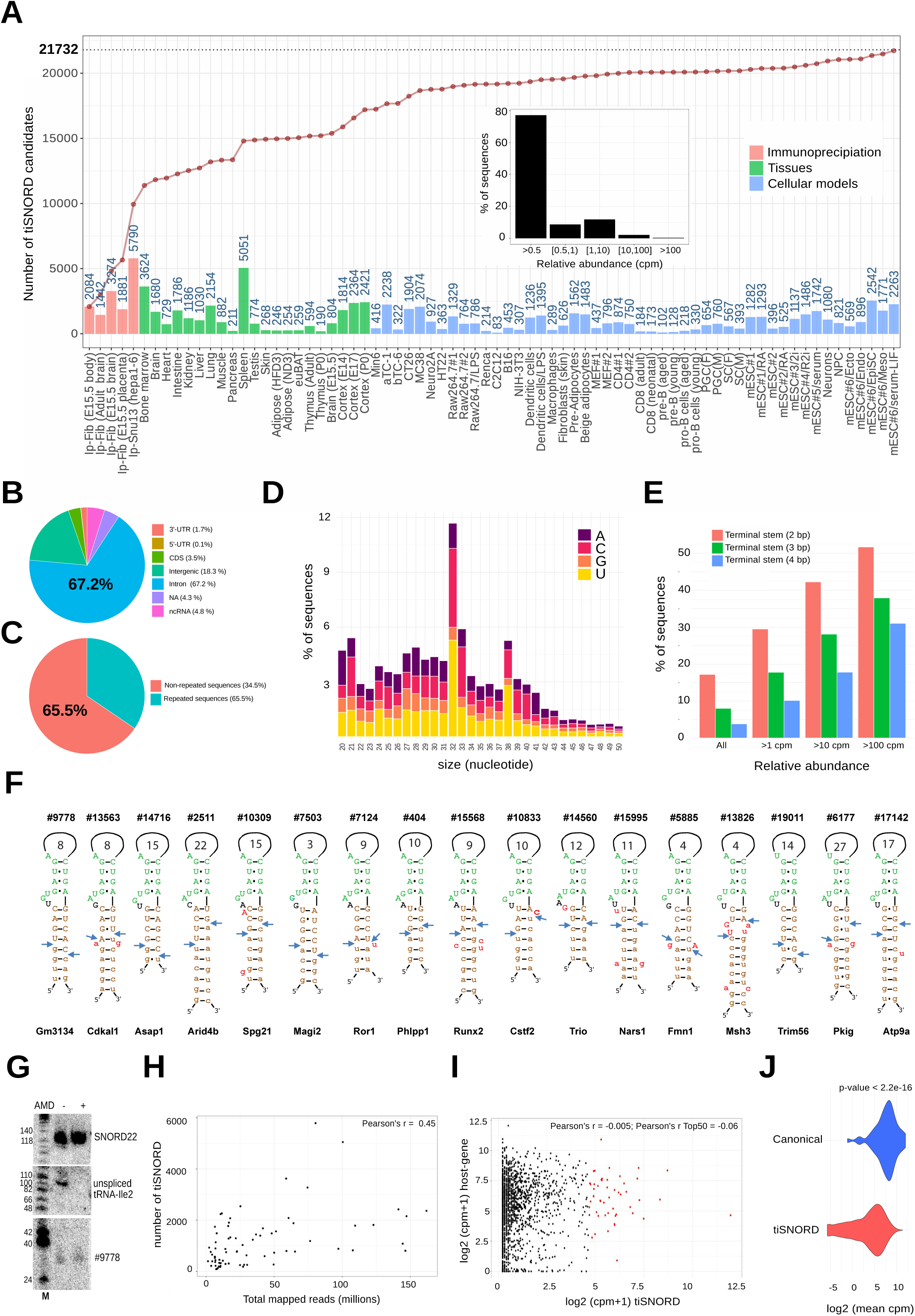
Identification of box C/D RNA-like sequences in publicly available small RNA-seq datasets. (A) Cumulative number of tiSNORDs detected across small RNA-seq datasets generated from IP experiments (Figures 1 and 2) or detected in mouse cells or tissues (Supplementary Table 3). The number of tiSNORDs identified in each biological sample is indicated above each bar. Inset: bars show the percentage of tiSNORDs based on their relative abundance, as measured by counts per million (cpm). (B) Pie chart showing the relative proportions (%) of tiSNORDs according to their genomic location. (C) Pie chart showing the relative proportions (%) of tiSNORDs derived from interspersed repetitive elements. (D) Size distribution (nt) of tiSNORDs, with the proportion of sequencing reads starting with A, C, G or U shown for each size class. (E) The prevalence of 5′-3′ terminal stem structures (2, 3, or 4 base pairs) correlates with tiSNORD abundance (F) Schematic representation of 5′-3′ terminal stem structures, including extended and irregular helices involving flanking intronic sequences (shown in lowercase). Nucleotide positions disrupting the canonical stem structure are shown in red while box C and box D motifs are indicated in green. Blue arrowheads mark the 5′ and 3′ termini of the fully processed SNORD candidates. The invariant unpaired nucleotide residue upstream of the C box is shown in black. The host gene is indicated below and the size of the spacer (nt) between the C- and D-boxes is shown within the loop. (G) Northern blot analysis of tiSNORD#9778 (15% denaturing PAGE) in undifferentiated mouse ES cells treated or not with AMD, as indicated above the panel. SNORD22 and tRNA-Ile2 were used as internal controls (stable and unstable RNAs, respectively). M: molecular markers (nt). (H) The scatter plot shows the relationship between the number of identified tiSNORDs and the sequencing depth of the biological samples analyzed. (I) The scatter plot shows the relationship between the relative expression of tiSNORDs and their host-gene transcripts, as measured in undifferentiated mES cells. The Top 50 most highly expressed tiSNORDs are denoted as red dots. (J) Expression levels of tiSNORD host gene transcripts are compared with those of canonical SNORD host gene transcripts, as measured in undifferentiated mES cells.

Using the complete set of tiSNORDs, we found that 67.2% are located within introns (Figure 3B) of at least ∼8,514 annotated transcripts, while 65.5 % do not overlap annotated interspersed repeats (Figure 3C). Unlike canonical SNORDs, intron-encoded tiSNORDs do not exhibit a preferential localization ∼70-80 nt upstream of the 3′ splice site (57) and are uniformly distributed across intronic regions (Supplementary Table 4). More importantly, they are generally weakly expressed (Figure 3A-inset), with only 12% represented at more than one count per million (cpm). Moreover, they appear to be poorly conserved; a detailed conservation analysis revealed that only 10% of the Top 100 most highly expressed murine tiSNORDs are conserved in *R. norvegicus* (not shown). Finally, the size distribution of tiSNORDs exhibits an apparent peak at 32 nt, accompanied by a pronounced preference for pyrimidine residues at the 5′ end, a feature that is also observed, at least to some extent, for some other lengths notably 33 nt and 38 nt (Figure 3D). The functional significance of this pattern, if any, remains unclear.

Many canonical SNORDs form a characteristic 4-5 bp terminal stem that leaves the nucleotide upstream of the C-box unpaired and sometimes involves additional flanking intronic sequences. This secondary structure facilitates k-turn folding and the post-transcriptional processing of SNORDs from their host gene introns, even though some SNORDs lacking such a terminal helix are nonetheless processed efficiently (58). Although not included as an essential criterion in our bioinformatic pipeline, we nevertheless evaluated whether the 5′ and 3′ ends of tiSNORDs have the potential to form base-pairing interactions. As indicated in Figure 3E, 17.2%, 8% and 3.8% of tiSNORDs may form a canonical 5’-3’ terminal stem structure comprising of at least 2, 3, and 4 contiguous base-pairs, respectively. This proportion is likely underestimated as manual inspection uncovered irregular RNA helices - including longer external intronic ones - not detectable computationally, as exemplified in Figure 3F. Remarkably, the relative positioning of the mature 5’ and 3’ ends of tiSNORDs is defined regardless of the length and regularity of the helix. This feature is another characteristic of the exonucleolytic post-transcriptional trimming of canonical intronic SNORDs (16, 58). In addition, consistent with their putative involvement in SNORD processing, stability and RNP assembly, the prevalence of such predicted terminal helices among tiSNORDs correlates well with their relative abundance, reaching 52.9%, 39.2% and 33% for 2-, 3-, and 4-bp helices, respectively, in the most abundant fraction (>100 cpm; Figure 3E).

Finally, we sought to detect endogenous expression of tiSNORD#9778, one of the most consistently identified SNORD candidates, notably in mouse embryonic stem (ES) cells (∼3500 cpm). As shown in Figure 3G, Northern blot analysis of 20 µg of total RNA from undifferentiated mES cells revealed an expected weak signal that, remarkably, persisted after 5 hours of actinomycin D treatment (5 µg/ml; AMD), a condition known to rapidly inhibit all RNA polymerase-dependent transcription. This observation indicated a steady-state stability of tiSNORD#9778 comparable to that of SNORD22, which was used as an internal control. On the contrary, and as anticipated for short-lived RNA intermediates, intron-containing tRNA-Ile2 signals were no longer detected in AMD-treated cells. This set of observations - presence of terminal stem and *in vivo* stability of endogenously-expressed tiSNORDs - reinforces our initial claim that tiSNORDs are structurally related to canonical SNORDs.

While their recurrent detection across independent datasets supports their authenticity, the overall low abundance of tiSNORDs, often represented by only a few sequencing reads, poses both technical and biological challenges. Although their detection might be expected to scale with the sequencing depth of the examined datasets, this relationship showed only a relatively weak tendency, implying that increasing sequencing depth in a given sample does not necessarily lead to a substantial increase in detected tiSNORDs (Figure 3H). Accordingly, tiSNORD repertoires show only partial overlap across small RNA-seq datasets when generated from the same cell type in independent studies, as illustrated for mES, mouse embryonic fibroblasts (MEFs), CD4 T cells and RAW 264.7 macrophages (Supplementary Figure 3A). Furthermore, although strong SNORD signatures were detected in 10 out of 11 mouse tissues studied by Isakova et al. (2020) (59), they were surprisingly largely absent from an independent small RNA-seq dataset covering 16 mouse tissues studied by Wagner *et al.* (2024) (60), as exemplified for adult mouse brain in Supplementary Figures 3B-F. Although we do not yet have an obvious explanation for these discrepancies, we posit that tiSNORD detection may be influenced by technical biases or artifacts arising during RNA extraction and/or library preparation. As a corollary, any direct comparison between two datasets generated by different studies - and thus probably different protocols of RNA extraction and library preparation - should be interpreted with extreme caution. Keeping these considerations in mind, and despite the absence of SNORD signatures in the bulk sequencing reads reported by Wagner et al. (2024), we nevertheless identified 4887 tiSNORDs, including 1640 that had not been previously reported (Supplementary Figure 3G). To avoid introducing false-positive signals, the sequences of these newly identified tiSNORDs are provided separately in Supplementary Table 5. Although additional tiSNORDs may emerge from the analysis of further biological samples and/or experimental conditions (*e.g.,* stress conditions), the apparent plateau observed in the cumulative discovery curve (Figure 3A; Supplementary Figure 3G) indicates that the catalog of the most highly expressed mouse tiSNORDs is likely approaching completion.

We next compared tiSNORD expression with that of their host genes, using undifferentiated mES cells for which both mRNA-seq and small RNA-seq datasets were generated in the same study (61). As illustrated in Figure 3I, tiSNORD abundance is not strictly dependent on expression level of their host-gene transcript, even when considering the top 50 most highly expressed tiSNORDs. Even more strikingly, tiSNORDs are preferentially hosted by relatively weakly expressed genes (Wilcoxon test, P < 2.2 × 10 ¹ ; Figure 3J), compared to canonical SNORDs which are mostly intron-encoded within ubiquitously expressed host genes encoding ribosomal proteins and translation factors. Although we did not identify any strictly tissue- or cell-specific tiSNORDs with expression levels exceeding 1 cpm, the use of Top 200 most highly expressed tiSNORDs is generally sufficient to infer bioinformatically the tissues of origin, as illustrated in Supplementary Figure 4, for a series of 11 mouse tissues. These findings not only provide a straightforward explanation for the inherently low abundance of tiSNORDs (*i.e.,* their low expression levels may be largely determined by those of their host genes), but also argue against the simplistic transcriptome background model according to which low-abundance tiSNORDs merely represent degradation by- products of highly expressed transcripts.

To investigate tiSNORD molecular biology, a selection of six tiSNORDs (tiSNORD#9778, #14716, #15995, #13826, #6177 and #2511), together with their flanking intronic sequences (Supplementary Data 1), were PCR-amplified and inserted into the second intron of the human β-globin gene, a well-established construct routinely used for ectopic expression of SNORDs (62). The resulting plasmids were transiently transfected into mouse L929 fibroblast cells and tiSNORD production was assessed by Northern blot analysis. As shown in Figure 4A, five out of six tested tiSNORDs produced a detectable signal whose size is consistent with that of the corresponding sequencing reads with, as expected, some heterogeneity in the apparent size of the detected signals. Primer extension analysis performed on tiSNORD#9778 and tiSNORD#14716, which exhibited the strongest Northern blot signals, demonstrated that their 5′ ends are correctly positioned 4-5 nucleotides upstream of the C-box (Figure 4B). Consistent with this, both tiSNORDs were specifically detected in anti-fibrillarin immunoprecipitates, indicating their incorporation into box C/D RNP particles (Figure 4C). We thus conclude that tiSNORDs can be reliably expressed from heterologous intronic contexts, as previously reported for canonical SNORDs. The weaker or apparent lack of detection for some tiSNORDs (#15995) may result from the limited sensitivity of Northern blotting and/or from improper folding of the inserted tiSNORDs in an ectopic sequence environment which could, through steric hindrance, prevent proper k-turn formation.

**Figure 4.**
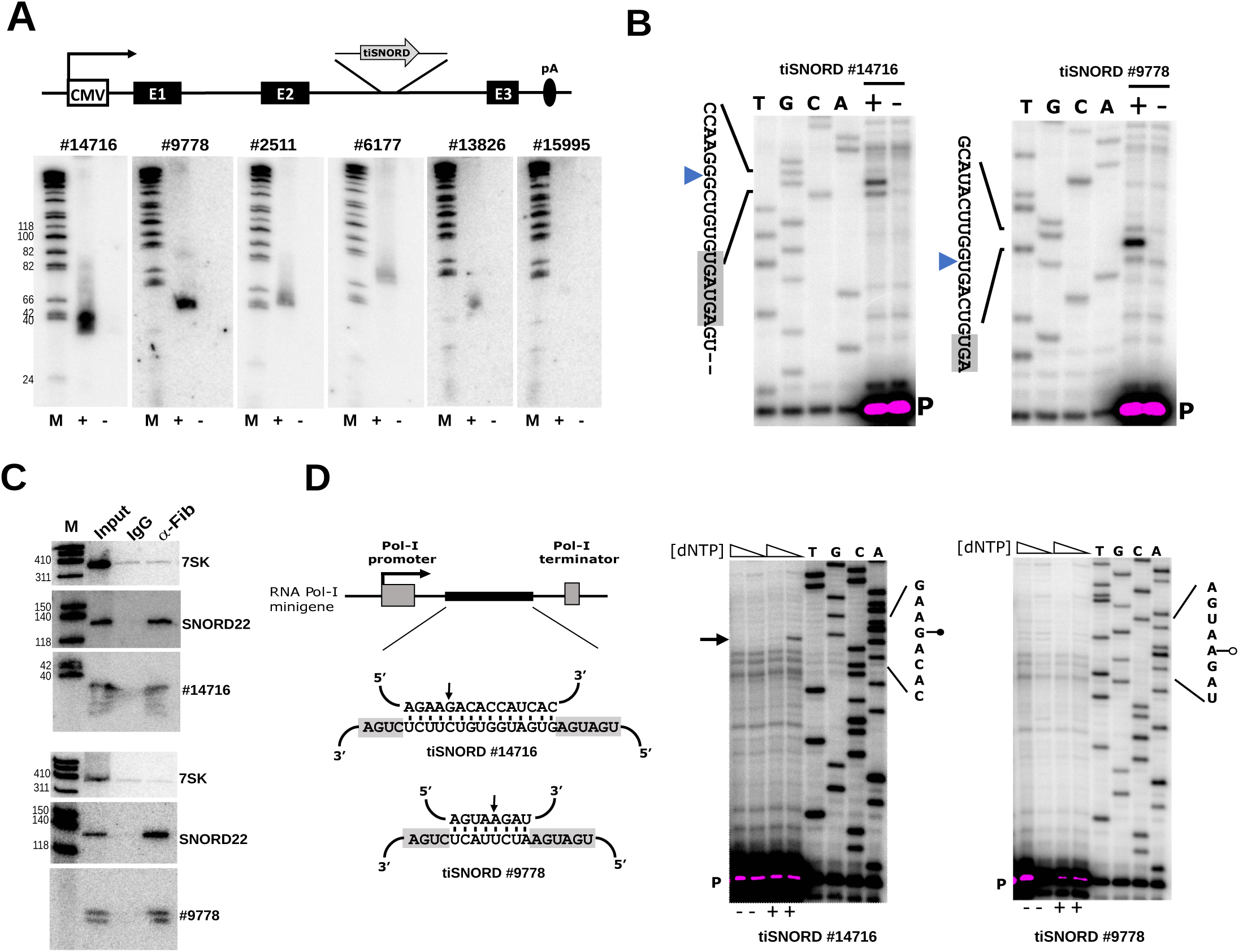
tiSNORDs retain their ability to target site-specific ribose methylation. (A) Top: Schematic representation of the β-globin minigene under the control of an RNA polymerase II (CMV) promoter. The sequences of inserted tiSNORD seqeunces are provided in Supplementary Data 1. Bottom: Northern blot analysis of tiSNORDs (15% denaturing PAGE), as indicated above the panel, in transiently transfected L929 cells (+) compared to non-transfected controls (-). M: molecular markers (nt) (B) 5′-end mapping of tiSNORD#14716 and tiSNORD#9778 by primer extension in transiently transfected L929 cells (+) compared to non-transfected controls (-). cDNAs were resolved onto 12% denaturing PAGE. The tiSNORD-expressing plasmids were sequenced in parallel and the ^32^P-labeled oligonucleotide is indicated as P. Blue arrowheads indicate the main detected 5’ termini. (C) Immunoprecipitation using anti-fibrillarin (72B9) antibody. tiSNORD#9778 and tiSNORD#14716 were detected by Northern blot (8% denaturing PAGE), with 7SK and SNORD22 serving as negative and positive controls, respectively. M: molecular markers (nt). (D) Left: Schematic representation of base-pairing interactions between a Pol I-synthesized substrate (top) and a tiSNORD (bottom). The C- and D- boxes are highlighted in grey and the nucleotide residue predicted to be targeted for ribose methylation according to the “+5 nt rule” is indicated by a small arrow. Note that the actual size of the RNA duplex is not known; only the longest theoretical duplex is shown. Right: Detection of ribose-methylated sites on RNA polymerase I transcripts following co-transfection (+) or no co-transfection (-) with tiSNORD#14716 or tiSNORD#9778, as indicated above each panel. Ribose methylation sites are identified by reverse transcription stops (black arrows), which occur one nucleotide before the modified position and are specifically observed under low dNTP concentration.

Despite their reduced size, we reasoned that tiSNORDs may have the ability to function as RNA methylation guides, provided that the spacer between the C and D boxes is sufficiently long to accommodate a 10-nt antisense sequence. Indeed, truncation experiments have previously shown that artificially truncated SNORDs, as short as 30-40 nt, can accumulate in the nucleolus and remain functionally competent (11, 15, 63). Moreover, structural and biochemical studies in Archaea have established that efficient *in vitro* ribose methylation requires a minimal spacer of 12 nt between the C- and D-boxes, together with an RNA duplex of 10-bp (64). These observations prompted us to investigate whether tiSNORD#14716, a 36-nt-long RNA with a 15-nucleotide spacer, is capable of guiding sequence-specific 2′-O-methylation.

To this end, tiSNORD#14716-expressing plasmids were co-transfected with an RNA polymerase I (Pol I) minigene construct expressing a 15-nt complementary target region (Figure 4D-left). As shown in Figure 4E-right, primer extension assays performed under both high- and low-dNTP conditions revealed a reverse transcriptase pause located one nucleotide downstream of the predicted target nucleotide, only under low-dNTP conditions. As expected, this RT signal was absent in control cells lacking tiSNORD#14716 expression. Of note, tiSNORD#9778, which contains an 8-nt spacer (Figure 4D-left), was unable to target the Pol I RNA substrate (Figure 4D-right), possibly due to a shortened and/or suboptimal spacer and antisense element. Although primer extension assays using low dNTP concentrations are not a quantitative assay (65), these experiments nevertheless demonstrate that at least some tiSNORDs may, in principle, retain the capacity to function as canonical nucleolar RNA methylation guides.

## Discussion

By combining anti-Fibrillarin and anti-Snu13 immunoprecipitation with systematic mining of a large collection of small RNA-seq datasets spanning diverse murine biological samples, we substantially expand the catalog of mouse box C/D small RNAs, uncovering an unexpectedly extensive repertoire that increases considerably the number of SNORDs from only a few hundred previously annotated loci to more than 20,000. However, although these newly identified SNORD candidates closely resemble canonical intronic SNORDs in terms of sequence features, processing pathways, RNP assembly and RNA-guided 2’-O-methylation activities, they differ markedly by their lower sequence conservation and reduced expression levels and, in the case of tiSNORDs, by their shorter length, ranging from 20 to 50 nt. Consistent with this latter characteristic, we noted that a small number of poorly conserved and low-abundance “mini box C/D RNA-like” molecules have previously been reported in HeLa cells (53) and may represent the human counterparts of mouse tiSNORDs. Even though we provide independent, complementary and unbiased evidence that SNORD candidates are not merely random transcriptomic byproducts, we acknowledge that establishing their evolutionary and biological relevance represents a distinct and challenging question that lies beyond the scope of the present study. Nevertheless, we consider it important to discuss several non-mutually exclusive hypotheses that could account for their existence, including their limited evolutionary conservation and low abundance, two features that could otherwise be interpreted as arguments against functionality.

In line with the notion that most, if not all, mouse and human rRNA 2′-O-methylation sites have already been mapped and convincingly assigned to canonical SNORDs (66, 67), our analyses did not identify any novel SNORDs whose expression and conservation levels were comparable to those of canonical SNORDs. Accordingly, it is unlikely that the vast majority of these newly identified SNORD candidates function as guides for robust, constitutive rRNA 2′-O-ribose methylations. Moreover, the very short spacing between the C and D boxes in many tiSNORDs is incompatible with the formation of an antisense element long enough to support specific RNA target recognition. Nevertheless, our proof-of-principle results using nucleolar Pol-I minigene demonstrate that certain tiSNORDs can direct sequence-specific ribose methylation, raising the possibility that at least a subset of them may target previously unrecognized RNA substrates, including possibly mRNAs. Consistent with this possibility, emergent studies have identified internal 2′-O-methylation sites within mRNAs that are typically only fractionally methylated (<20-30%), suggesting the existence of tightly regulated and still poorly understood targeting mechanisms (68–72). Assuming a role in targeting mRNAs for methylation, the low abundance of tiSNORDs may reflect highly restricted temporal or cell type-specific expression patterns.

Beyond a role in RNA modification, additional possibilities must also be considered, including the most parsimonious one, namely biological noise. The number of sequences capable of forming a k-turn-like structure in the mouse transcriptome is likely far greater than the number of *bona fide* SNORDs identified to date, and how intronic SNORD sequences are specifically recognized by Snu13 and other assembly factors is not entirely understood (10). Accordingly, one straightforward explanation is that numerous tiSNORDs are mistakenly recognized as a result of stochastic errors during RNP assembly. In a more extreme scenario, one could even speculate that many, if not all cryptic k-turns in a given transcriptome are transiently surveyed by the SNORD biogenesis machinery, with the resulting processed k-turn-containing RNAs persisting in the cell without being subject to strong selective pressure. Accordingly, our findings may primarily reflect the sensitivity of NGS-based approaches in revealing the plasticity and error-prone molecular processes underlying box C/D RNP assembly rather than the discovery of novel RNA-mediated regulatory functions. Evolution of mammalian SNORDs is an ongoing process involving birth and death in a lineage or even species-specific manner (73–75). From long terms evolutionary perspectives, accidental production of SNORD-like RNAs could become beneficial as long as they reach higher expression levels and acquire antisense element conferring cellular fitness and selective advantages. In other words, SNORD candidates could also be viewed as continuously-produced raw substrates from which novel SNORDs might emerge during the course of evolution. In this line of thought, that one third of SNORD candidates originate from interspersed repetitive elements raises the possibility that “genomic parasites” can be co-opted into regulatory small RNAs, as already well established for the murine, brain-specific BC1 RNA (76). The case of the primate-specific, weakly-expressed Alu-ACA RNAs originating from Alu elements represents another example of recent “domestication” of repeated elements in snoRNA-like RNA species (77).

Assuming that Snu13 has the potential to interact with any encountered k-turn, thousands of transient - yet specific - interactions may also contribute to confining a pool of Snu13 within subnuclear compartments where pre-mRNA splicing and RNP assembly may preferentially take place. By locally increasing its concentration, this could, in turn, facilitate the assembly of canonical box C/D RNPs. This scenario, which implies a role in nuclear organization, is consistent with models proposing that networks of weak interactions modulate the intranuclear diffusion of transcription factors, thereby influencing their ability to efficiently locate and recognize their cognate DNA binding sites (78). An alternative, yet related, interpretation is that these newly-identified, unprocessed k-turn motifs recruit, and potentially sequester, box C/D core RNA binding proteins in a context-dependent manner. Although we consider such a mechanism unlikely, at least under physiological stoichiometric conditions examined here, the ncRNA LoNA harboring a C/D box-like motif has been proposed to function as an RNA decoy that modulates the activity of Fibrillarin and Nucleolin (79). Another possibility is that early recruitment of box C/D RNP factors onto k-turns affects - directly or indirectly - the overall intra-nuclear fate of the host-gene transcript (*i.e.,* its release from chromatin environment, its splicing or its packaging in mRNPs).

Consistent with this idea, unprocessed form of SNORD86 adopts different RNP conformations and influences alternative splicing of its host Nop56 pre-mRNA (80). Of note, box C/D RNA-like element embedded within the ncRNA *CRNDE* has recently been shown to play some roles in proliferation and ribosome biogenesis of renal cell carcinoma (81). Finally, SNORD candidates may participate in stabilizing some yet-to-be-discovered snoRNA-retaining transcripts (82) or long noncoding RNAs with snoRNA-derived ends (83). Thus, the above-mentioned hypotheses posit that some SNORD candidates, and especially their k-turn may primarily function as *cis*-regulatory elements (*i.e.,* k-turn serving as RNA-binding sites) without undergoing efficient or constitutive processing, which could account for their intrinsically low abundance. Under this assumption, sequence conservation beyond the k-turn may not be strictly required, provided that binding of Snu13 is preserved.

The k-turn is a recurrent and ancient architectural RNA motif found across a wide range of functionally and structurally diverse RNA classes, including ribosomal RNA, spliceosomal U4 snRNA, riboswitches, yeast mRNAs, RNase P, SRP RNA, and archaeal box C/D and H/ACA small RNAs (84, 85). This widespread occurrence across RNA molecules is consistent with an “RNA bricolage model”, in which a structural RNA motif has been repurposed multiple times to fulfill diverse functions. Accordingly, SNORD candidates - and tiSNORDs in particular - may constitute a previously unrecognized class of k-turn-containing small RNA species. Each tiSNORD would be an individually low-abundance and poorly conserved RNA molecule, yet they could collectively contribute to cellular processes. A parallel can be drawn, at least to some extent, with pre-pachytene piRNAs, which also comprise a huge population of individually rapidly evolving, often low-abundance ∼ 26-31 nt-long small RNAs that nonetheless act in mass to ensure genome defense (86). Under this appealing, albeit highly speculative hypothesis, tiSNORDs may exert yet-to-be-discovered *trans*-acting regulatory functions that are not necessarily mediated by base-pairing interactions.

In conclusion, our findings provide a valuable resource for future investigations into this unexpectedly large and previously unrecognized family of box C/D RNA-like species. More sophisticated molecular and mechanistic studies, together with comprehensive cross-species analyses across diverse organisms, will be required to test the hypotheses outlined above and fully assess the biological and evolutionary significance of our findings. A key question remains: do tiSNORDs and the other SNORD candidates identified here represent the visible tip of an iceberg (*e.g.,* a novel class of box C/D-like RNA species acting through *cis*- and/or *trans*-regulatory mechanisms) or do they instead reside in the abyss of transcriptomes as by-products of SNORD processing pathways (*e.g.,* biological noise with limited functional relevance)?

## Supplementary Figures - Legends

**Supplementary Table 1** - This file lists the number of sequencing reads mapping to annotated non-coding RNAs (Supplementary Table 6) in the four mouse tissues analyzed by anti-Fibrillarin IP, as well as in the Hepa 1-6 cell line analyzed by anti-Snu13 IP.

**Supplementary Table 2** - This file lists the number of sequencing reads corresponding to SNORD candidates identified in the first and second screening steps in the four mouse tissues analyzed by anti-Fibrillarin IP, as well as in the Hepa 1.6 cell line analyzed by anti-Snu13 IP.

**Supplementary Figure 1** - Identification of novel box C/D RNA-like RNAs in developing (E15.5) brain, body and placenta (First screen). (A) Distribution of the positions of the C-box relative to the 5′ end. (B) Distribution of the positions of the D-box relative to the 3′ end. (C) Relative proportions of C-box sequence variants. (D) Relative proportions of D-box sequence variants. (E) Overrepresentation of uridine residues upstream of the D-box in selected SNORD candidates (second screen). The dashed red line indicates 25%, the proportion expected by random sampling.

**Supplementary Table 3** - This file lists the accession numbers of all RNA-seq datasets analyzed in this study and retrieved from the Sequence Read Archive (SRA) database (https://www.ncbi.nlm.nih.gov/sra).

**Supplementary Table 4 -** This file provides the complete catalog of SNORD candidates, including both tiSNORDs (20-50 nt) and other SNORD candidates (>50 nt), identified from anti-fibrillarin and Snu13 immunoprecipitation experiments and from publicly available small RNA-seq datasets

**Supplementary Figure 2** - Identification of novel box C/D RNA-like RNAs in small RNA-seq datasets generated from bone marrow, undifferentiated ES cells and α-TC1 pancreatic cell lines (First screen). (A) Distribution of the positions of the C-box relative to the 5′ end. (B) Distribution of the positions of the D-box relative to the 3′ end. (C) Relative proportions of C-box sequence variants. (D) Relative proportions of D-box sequence variants. (E) Overrepresentation of uridine residues upstream of the D-box (second screen). The dashed red line indicates 25%, the proportion expected by random sampling.

**Supplementary Figure 3 -** Experimental limitations of computational tiSNORD detection. (A) tiSNORD repertoires show only partial overlap across different small RNA-seq datasets generated from the same cell type, as illustrated for CD4 cells, mES cells (datasets #1-#5), RAW 264.7 macrophages and MEFs, and also to some extent between biological replicates (mES#6; Rep1-Rep3). (B-F) Mouse brain small RNA-seq datasets generated by Isakova et al., (2020) but not those generated by Wagner et al., (2024) display overt canonical SNORD molecular signatures. The same holds true for the other analyzed tissues (not shown). (B) Distribution of the positions of the C-box relative to the 5′ end. (C) Distribution of the positions of the D-box relative to the 3′ end. (D) Relative proportions of C-box sequence variants. (E) Relative proportions of D-box sequence variants. (F) Overrepresentation of uridine residues upstream of the D-box. The dashed red line indicates 25%, the proportion expected by random sampling. (G) Cumulative number of novel tiSNORDs detected across 16 small RNA-seq datasets of mouse tissues (old and young mice) generated by Wagner *et al.*, (2024) The number of tiSNORDs identified in each biological sample is indicated above each bar (see also Supplementary Table 5).

**Supplementary Table 5 -** This file provides the catalog of novel tiSNORDs identified in small RNA-seq datasets from 16 mouse tissues (aged and young) generated by Wagner *et al*. (2024).

**Supplementary Figure 4 -** The Top 200 highly expressed tiSNORD host-gene transcripts enable identification of their tissue of origin, as assessed using Mouse Gene Atlas analyses (87).

**Supplementary Data 1** - Genomic sequences cloned into the intron of the β-globin gene, with tiSNORD sequences shown in bold and the C and D boxes highlighted in green.

**Supplementary Table 6** - Sequences of non-coding RNAs retrieved from GtRNAdb, RNAcentral, SnoPY, Ensembl and miRBase, used to filter the small RNA-seq datasets.

**Supplementary Table 7** - Sequences of DNA oligonucleotides used in this study.

## Methods

### Tissue extracts, immunoprecipitation and Illumina library preparation

Upon euthanasia, mouse tissues were harvested, snap-frozen immediately in liquid nitrogen, and stored at -80 °C. Tissues were disrupted at 4 °C using an Ultra-Turrax homogenizer in NET-150 buffer (50 mM Tris pH 7.4, 150 mM NaCl, 0.05% Igepal CA-630 (Sigma), supplemented with Promega RNasin Ribonuclease Inhibitor and Roche protease inhibitors. Tissue extracts were clarified by centrifugation (10,000 rpm in a Sorvall SS34 rotor for 10 min at 4°C), and the supernatant was subjected to immunoprecipitation using anti-fibrillarin (72B9) antibodies (a gift from M. Pollard). Briefly, 100 µl of hybridoma 72B9 supernatant was incubated for 120 min at 4 °C with 80 µL of Protein A Sepharose® 4B Fast Flow (S. aureus; Sigma #P9424) in 1 ml NET-150. Beads were then washed four times with 1 ml NET-150 and subsequently incubated with 1 ml of tissue extract for 60 min at 4 °C under gentle agitation. Immunoprecipitates were collected by centrifugation and washed seven times with 1 ml NET-150. RNA was extracted from the pellet by SDS/phenol extraction. cDNA libraries were prepared using the Illumina TruSeq Small RNA Library Prep Kit and sequenced on an Illumina HiSeq platform in a paired-end 2×150 bp configuration. Library preparation and next-generation sequencing were performed by Genewiz (https://www.genewiz.com/en-GB/). Raw next-generation sequencing data have been deposited in the Gene Expression Omnibus (GEO) under accession number GSE333474.

### Computational search for box C/D RNA-like sequences

Raw FASTQ files were quality-checked using FastQC (v0.12.1) to assess sequencing quality and adapter contamination (https://www.bioinformatics.babraham.ac.uk/projects/fastqc/). Although some libraries were sequenced in paired-end mode, only R1 reads were retained for downstream analyses, as small RNA-seq workflows rely on short insert sizes and the information contained in R2 is typically redundant or dominated by adapter sequence. Moreover, retaining R2 reads rarely improves mapping or annotation of short RNAs and introduce unnecessary complexity when all relevant information is already captured in R1. Adapter removal, optional UMI trimming and quality filtering were performed using Trim Galore (v0.6.10), which calls Cutadapt (v5.1) (https://www.trimgalore.com/; DOI:10.14806/ej.17.1.200). Cutadapt parameters included a maximum error rate of 0.1 (-e 0.1), quality trimming at Phred score 20 (-q 20), a minimum adapter overlap of 1 (-O 1), and removal of reads shorter than 18 nt after trimming. Identical sequences were collapsed using custom AWK scripts and only sequences observed at least twice (≥2) were retained for downstream analyses. Filtered sequences were first compared against a manually curated collection of ncRNA sequences retrieved from GtRNAdb, RNAcentral, SnoPY, Ensembl, and miRBase (Supplementary Table 6) using BLAST (v2.14.0), with a minimum word size of 18 nt and an E-value cutoff of 0.05. (88). Note that canonical SNORD counting included both full-length and processed RNA species; however, only full-length species were considered when comparing canonical SNORDs with tiSNORDs. Furthermore, to minimize bias, the numerous tandemly repeated imprinted SNORDs (which account for one-third to one-half of annotated SNORDs) were grouped into three main families (SNORD116, SNORD115, and SNORD113/114), and a single randomly selected member from each family was retained for the analysis. Reads that did not match previously annotated ncRNAs were subsequently aligned to the mouse genome (mm10) using Bowtie2 (v2.5.1) in very-sensitive mode, allowing one mismatch in the seed (--very-sensitive -N 1) (89). All analyses and annotations were performed in R (v4.4.2) using a standard set of data-handling libraries (tidyverse, GenomicRanges, Biostrings) (90). Reads classified as known ncRNAs (BLAST hits) and non-BLAST reads mapping to mm10 were processed in parallel using the same annotation pipeline. Candidate mapped reads were annotated using annotatr (v1.32.0) with UCSC mm10 genomic annotations. Additional resources included BSgenome (v1.74.0), TxDb.Mmusculus.UCSC.mm10.knownGene (v3.10.0) and org.Mm.eg.db (v3.20.0), enabling assignment of promoter, exon, intron and intergenic categories, as well as gene identifiers. Overlap with repetitive elements was assessed using RepeatMasker annotations from UCSC (Smit, AFA, Hubley, R & Green, P. RepeatMasker Open-4.0.2013-2015 <http://www.repeatmasker.org) (91). Overlapping sequences were grouped into collapsed sequence clusters, and the most abundant sequence (“majority read”) was selected as the representative. The genomic sequence corresponding to each majority read was retrieved and scanned for putative C-box and D-box motifs (and variants) within windows anchored near the 5′ (C box, within 10 nt) and 3′ (D box, within 10 nt) ends of the sequence (1^st^ screen). When multiple candidate motifs were identified, a single best match was retained based on positional criteria (reference positions: 5 for C boxes and 2 for D boxes). A more stringent “selected set” (second screen) required the C box to occur at positions 4 or 5, the D box at positions 2 to 5, the C-box variant to belong to the set {C0, C1, C3, C4} and the D box to match the canonical D0 motif. From the selected pool, sequences were further stratified into “Selected tiny” ranging from 20 to 50 nt and “Selected long” (>50 nt). Upon manual inspection, we identified a small number of reads corresponding to processed forms of canonical SNORDs that had escaped filtering: these cases were therefore manually removed. Counts corresponding to candidate SNORDs were normalized to counts per million (cpm). The library size used for cpm calculation was defined as the total number of reads mapping to known ncRNAs plus the reads mapped to candidate RNAs. Sequences were further grouped based on at least 1-nt genomic overlap to identify redundant sequences across samples, enabling consistent cross-sample comparisons and assessment of conservation or presence/absence patterns. The full list of publicly available sRNA-seq datasets processed with the same pipeline is provided in Supplementary Table 3. All scripts and pipeline code are publicly available on GitHub: https://src.koda.cnrs.fr/biga_cbi/partageprojet/shared_projects/tisnord.

### Cell culture, Plasmids, Transfection

L929 cells (ATCC # CCL-1) were grown at 37°C with 5% CO2 in Dulbecco’s Modifier Eagle Medium (Gibco; 4.5 g/l glucose) supplemented with 1 mM Sodium Pyruvate (Gibco), 1% penicillin-streptomycin (Sigma-Aldrich) and 10% fetal bovine serum (PAN biotech). Transient transfection experiments were performed using JetPrime transfection reagent according to the manufacturer’s instructions. For one well of a 6-well plate (seeded the previous day with 400 000 cells), 1 µg of plasmid DNA (prepared using a Qiagen Maxiprep kit) was transfected using 200 µl JetPrime buffer and 4 µl of JetPrime reagent. Standard molecular biology techniques were used for DNA manipulation and cloning following established protocols. tiSNORD sequences together with their flanking intronic regions (Supplementary Data 1) were PCR-amplified and subsequently cloned into the ClaI-XhoI sites within the second intron of the human β-globin gene construct (62). Artificial Pol I target substrates were generated by inserting short complementary sequences, formed by annealed two oligonucleotides, into a Pol I minigene construct linearized with XbaI and XhoI. To stimulate Pol-I minigene expression, 24 h post-transfection cells were treated with a low concentration of actinomycin D (AMD, 0.05 µg/ml) for 120 min. The medium was then removed, cells were briefly washed with PBS and fresh medium (without actinomycin D) was added for an additional 36 h prior to RNA extraction (92).

### RNA extraction, Northern-blot, Primer extension

Total RNA was extracted from L929 cells using Tri-Reagent (Euromedex) following the manufacturer’s instructions. The RNA samples were subsequently treated with RNase-free DNase I (RQ1, Promega) and proteinase K (Sigma) to remove contaminating DNA and proteins. RNA was then resuspended in RNase-free water and stored at -20 °C until further use. For Northern blot analyses, 10-20 µg of total cellular RNA was separated by electrophoresis on 8%, 12% or 15% denaturing polyacrylamide gels (PAGE) containing 7 M urea. A molecular Marker was used (Promega ΦX174 DNA/HinfI Markers. RNA was then electroblotted onto a nylon membrane (Amersham Hybond-N, GE Healthcare) and UV-crosslinked using a Stratalinker. Hybridization was performed with 5′-end ^32^P-labeled DNA. Membranes were incubated overnight at 50 °C in hybridization buffer consisting of 5× SSPE, 5× Denhardt’s solution, 1% SDS, and 150 µg/mL yeast tRNA. Subsequently, membranes were washed twice for 15 min at room temperature in 0.1× SSPE/0.1% SDS. Radioactive signals were detected using a Typhoon™ Biomolecular Imager (Amersham). For mapping the 5′ ends of tiSNORDs, 5 µg of total RNA was mixed with a specific primer and heat-denatured for 1 min at 95°C, after which samples were allowed to cool to room temperature. Primer extension was then performed using AMV reverse transcriptase (Promega) at 42 °C for 60 min. For detection of 2′-O-methylation, the same primer extension procedure was applied, except that two dNTP concentrations were used: a high concentration (1 mM) and a low concentration (0.01 mM) known to induces a RT pause one nucleotide upstream of the ribose-methylated residue (65). DNA sequencing ladder was generated by primer extension using the same conditions with dideoxynucleotides elongation mixes. The sequences of the DNA oligonucleotides are provided in Supplementary Table 7.

## Acknowledgments

We thank M. Scott (University of Sherbrooke) and her colleagues for fruitful discussions throughout this work, as well as the staff of the BigA-T facilities at CBI (Toulouse University), particularly M. Aguirrebengoa and V. Rocher. We are also grateful to S. Labialle (IMoPA-CNRS, Nancy) for the careful and critical reading of the manuscript.

## CRediT authorship contribution statement

**Tomas Caetano:** review & editing, visualization, investigation, formal analysis, conceptualization. **Baptiste Grand:** review & editing, investigation, formal analysis; **Jade Hebras:** review & editing, investigation, formal analysis; **Michelle Feat:** investigation; **Matthias Zytnicki:** review & editing, visualization, investigation, formal analysis, conceptualization, supervision; **Jérôme Cavaillé:** writing - review & editing, investigation, formal analysis, conceptualization, supervision, project administration, funding acquisition.

## Author Declaration

All authors have approved the final version of the manuscript and confirm that it is not under consideration for publication elsewhere. The authors declare no conflicts of interest.

## Declaration of generative AI-assisted technologies

The authors used OpenAI’s ChatGPT solely to correct grammatical errors and improve the clarity of the English language. The authors reviewed and edited all AI-assisted text and take full responsibility for the final content. No AI tools were used to conduct bibliography searches, generate scientific ideas or hypotheses, interpret data or draw conclusions.

