## Supplementary figures and images for "A novel class of tiny box C/D-like RNAs reveals hidden complexity in the mouse box C/D RNA repertoire"

### Supplementary Figure 1

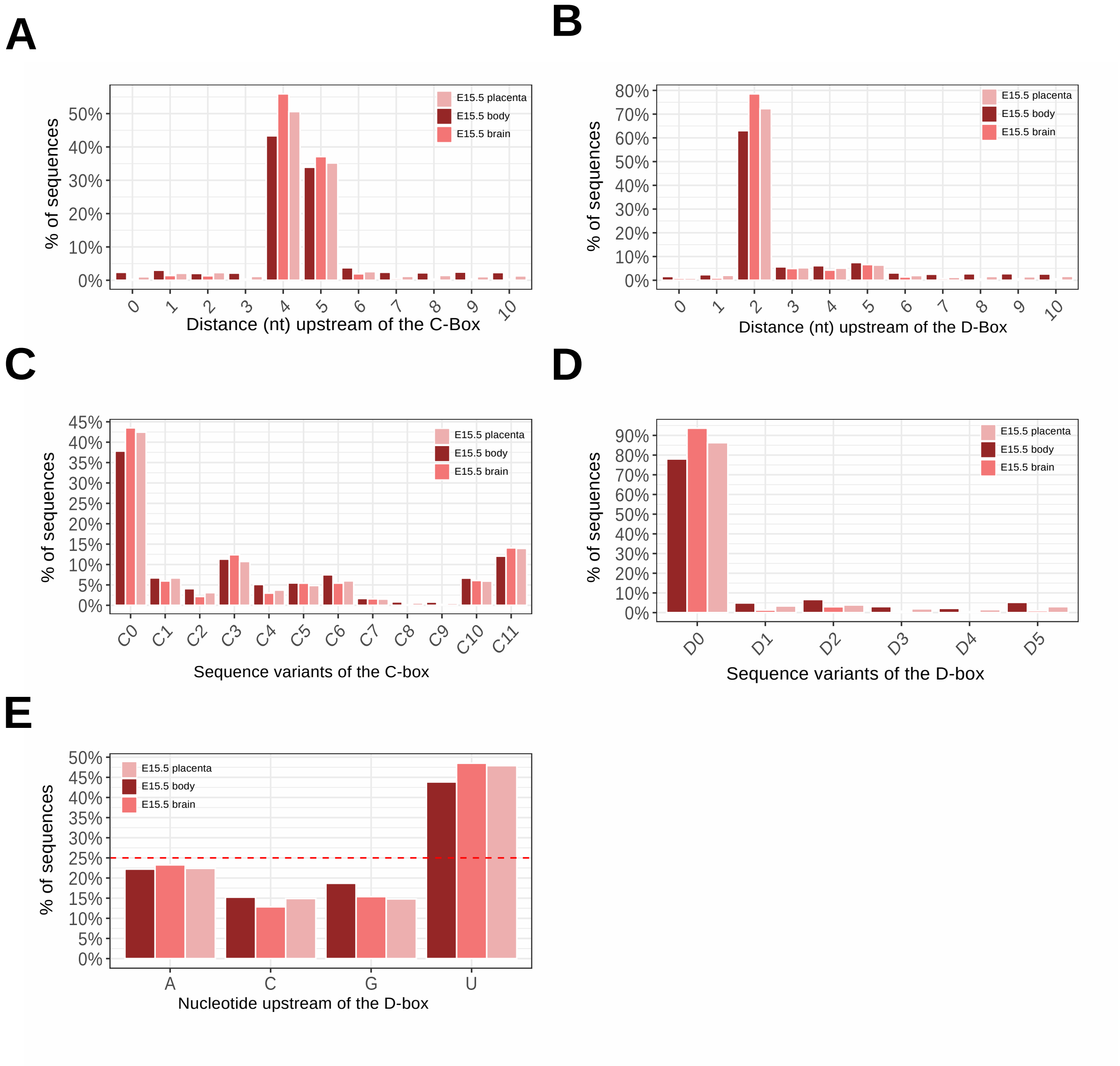


**Supplementary Figure 1 - Caetano et al**

### Supplementary Figure 2

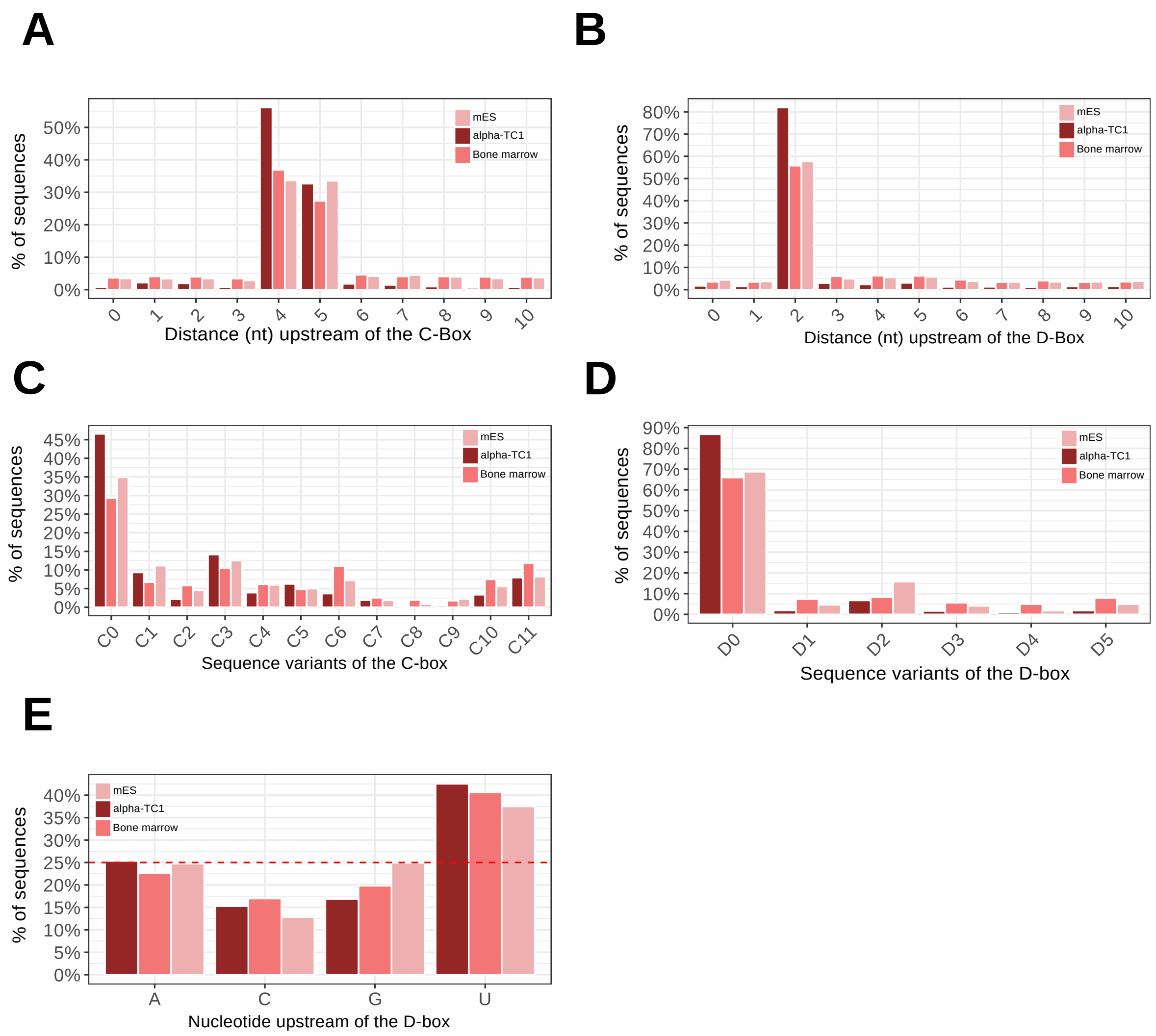


**Supplementary Figure 2 - Caetano et al**

### Supplementary Figure 3

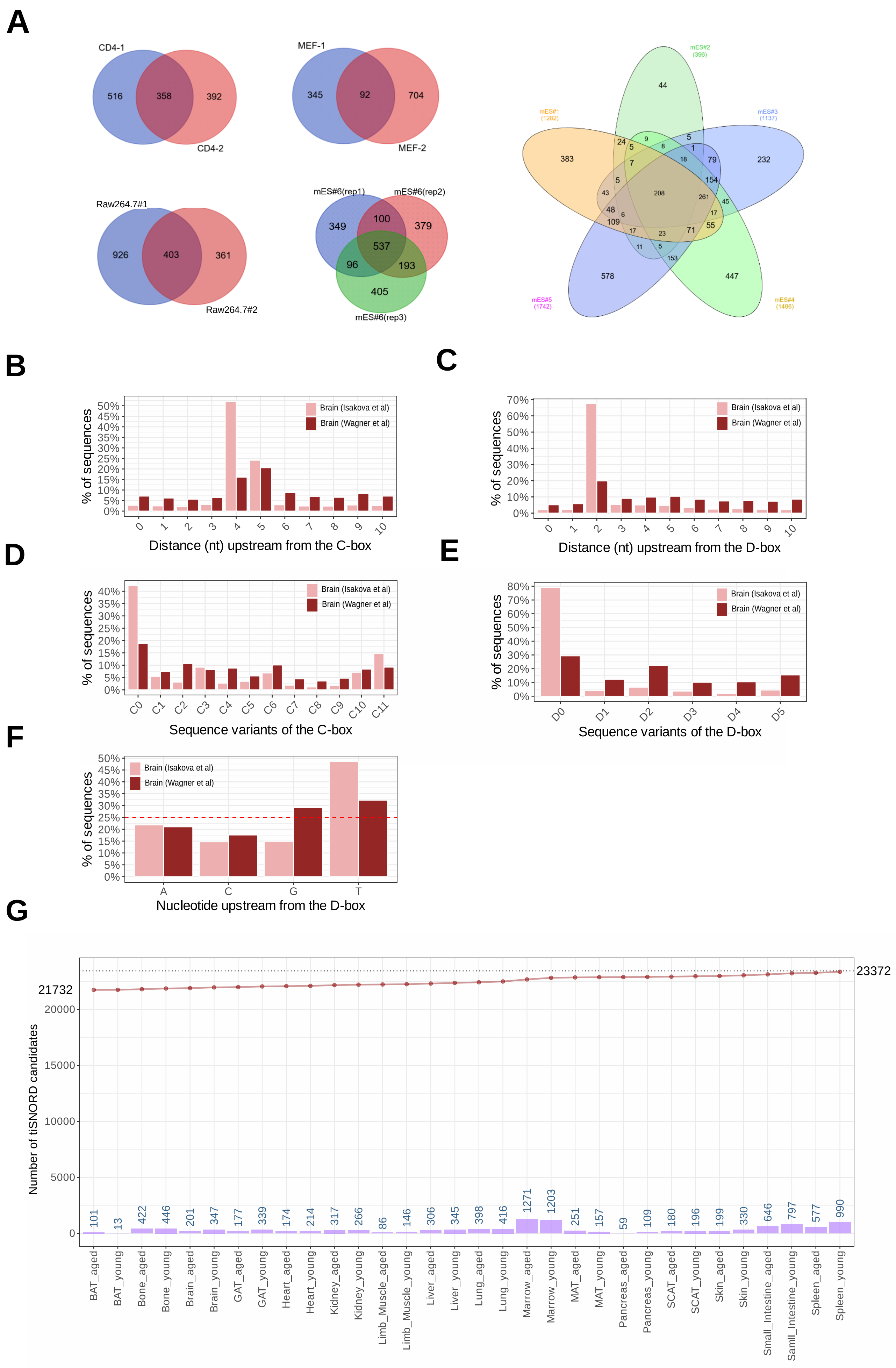


**Supplementary Figure 3 - Caetano et al**
