## Supplementary Figure 4 for "A novel class of tiny box C/D-like RNAs reveals hidden complexity in the mouse box C/D RNA repertoire"

### Top200 highly expressed (bone marrow)

| Index | Name | P-value | Adjusted p-value | Odds Ratio | Combined score |
|---|---|---|---|---|---|
| 1 | NK cells | 0.0001556 | 0.01261 | 2.99 | 26.23 |
| 2 | bone marrow | 0.005034 | 0.2039 | 2.77 | 14.65 |
| 3 | nih 3T3 | 0.008913 | 0.2406 | 3.16 | 14.91 |
| 4 | thymocyte DP CD4+CD8+ | 0.01388 | 0.2810 | 2.49 | 10.65 |
| 5 | mast cells IgE | 0.02282 | 0.3657 | 2.08 | 7.87 |
| 6 | stem cells HSC | 0.02897 | 0.3657 | 3.03 | 10.73 |
| 7 | skeletal muscle | 0.03160 | 0.3657 | 1.91 | 6.60 |
| 8 | follicular B-cells | 0.05264 | 0.5330 | 1.86 | 5.49 |
| 9 | neuro2a | 0.09831 | 0.8847 | 1.84 | 4.26 |
| 10 | Baf3 | 0.1124 | 0.9108 | 1.77 | 3.87 |

### Top200 highly expressed (brain)

| Index | Name | P-value | Adjusted p-value | Odds Ratio | Combined score |
|---|---|---|---|---|---|
| 1 | cerebellum | 9.272e-13 | 6.768e-11 | 7.83 | 216.84 |
| 2 | cerebral cortex | 9.912e-12 | 3.618e-10 | 8.23 | 208.65 |
| 3 | nucleus accumbens | 1.814e-8 | 4.415e-7 | 6.15 | 109.66 |
| 4 | dorsal striatum | 4.199e-7 | 0.000007664 | 4.87 | 71.52 |
| 5 | cerebral cortex prefrontal | 0.00002797 | 0.0004084 | 4.64 | 48.63 |
| 6 | olfactory bulb | 0.0001632 | 0.001985 | 3.82 | 33.28 |
| 7 | mast cells IgE | 0.003894 | 0.04061 | 2.59 | 14.39 |
| 8 | retina | 0.008722 | 0.07959 | 2.32 | 11.01 |
| 9 | amygdala | 0.01375 | 0.1103 | 2.89 | 12.40 |
| 10 | hippocampus | 0.01511 | 0.1103 | 3.62 | 15.16 |

### Top200 highly expressed (heart)

| Index | Name | P-value | Adjusted p-value | Odds Ratio | Combined score |
|---|---|---|---|---|---|
| 1 | heart | 1.849e-8 | 0.000001553 | 4.94 | 87.87 |
| 2 | skeletal muscle | 0.0004027 | 0.01691 | 2.85 | 22.25 |
| 3 | follicular B-cells | 0.0006984 | 0.01956 | 2.91 | 21.17 |
| 4 | bladder | 0.001469 | 0.03085 | 4.46 | 29.09 |
| 5 | cerebral cortex | 0.003182 | 0.05345 | 3.18 | 18.31 |
| 6 | umbilical cord | 0.009076 | 0.1271 | 3.53 | 16.61 |
| 7 | mast cells IgE | 0.01400 | 0.1680 | 2.25 | 9.62 |
| 8 | retinal pigment epithelium | 0.05087 | 0.5341 | 2.04 | 6.09 |
| 9 | NK cells | 0.06188 | 0.5425 | 1.81 | 5.03 |
| 10 | hippocampus | 0.06460 | 0.5425 | 2.74 | 7.50 |

### Top200 highly expressed (Intestine)

| Index | Name | P-value | Adjusted p-value | Odds Ratio | Combined score |
|---|---|---|---|---|---|
| 1 | intestine large | 0.007421 | 0.3728 | 2.77 | 13.60 |
| 2 | mast cells IgE | 0.008771 | 0.3728 | 2.32 | 10.97 |
| 3 | intestine small | 0.02716 | 0.5433 | 2.20 | 7.95 |
| 4 | skeletal muscle | 0.02939 | 0.5433 | 1.93 | 6.82 |
| 5 | salivary gland | 0.03196 | 0.5433 | 2.14 | 7.36 |
| 6 | NK cells | 0.04272 | 0.6051 | 1.87 | 5.91 |
| 7 | mIMCD-3 | 0.05691 | 0.6911 | 2.26 | 6.49 |
| 8 | thymocyte DP CD4+CD8+ | 0.08152 | 0.7722 | 1.93 | 4.83 |
| 9 | bone marrow | 0.08396 | 0.7722 | 1.91 | 4.74 |
| 10 | mast cells | 0.09903 | 0.7722 | 1.75 | 4.05 |

### Top200 highly expressed (kidney)

| Index | Name | P-value | Adjusted p-value | Odds Ratio | Combined score |
|---|---|---|---|---|---|
| 1 | kidney | 0.00001076 | 0.0008393 | 3.96 | 45.29 |
| 2 | mast cells IgE | 0.0001015 | 0.003960 | 3.39 | 31.20 |
| 3 | mast cells IgE+antigen 1hr | 0.02055 | 0.5342 | 2.93 | 11.37 |
| 4 | mast cells | 0.05708 | 0.7442 | 1.99 | 5.71 |
| 5 | cerebral cortex | 0.06347 | 0.7442 | 2.20 | 6.07 |
| 6 | macrophage bone marrow 6hr LPS | 0.07113 | 0.7442 | 1.76 | 4.65 |
| 7 | adipose brown | 0.07612 | 0.7442 | 1.96 | 5.06 |
| 8 | nucleus accumbens | 0.08973 | 0.7442 | 2.00 | 4.82 |
| 9 | NK cells | 0.09174 | 0.7442 | 1.72 | 4.11 |
| 10 | mast cells IgE+antigen 6hr | 0.1138 | 0.7442 | 2.62 | 5.68 |

### Top200 highly expressed (liver)

| Index | Name | P-value | Adjusted p-value | Odds Ratio | Combined score |
|---|---|---|---|---|---|
| 1 | liver | 7.435e-9 | 5.874e-7 | 4.06 | 76.01 |
| 2 | mast cells IgE | 0.0002256 | 0.008912 | 3.13 | 26.31 |
| 3 | adipose brown | 0.01799 | 0.4108 | 2.38 | 9.56 |
| 4 | skeletal muscle | 0.02080 | 0.4108 | 2.04 | 7.91 |
| 5 | macrophage peri LPS thio 0hrs | 0.03422 | 0.5407 | 2.37 | 8.01 |
| 6 | follicular B-cells | 0.07907 | 0.9695 | 1.78 | 4.51 |
| 7 | cerebellum | 0.1674 | 0.9695 | 1.65 | 2.95 |
| 8 | nih 3T3 | 0.1750 | 0.9695 | 1.87 | 3.26 |
| 9 | cerebral cortex | 0.1857 | 0.9695 | 1.69 | 2.84 |
| 10 | mast cells IgE+antigen 1hr | 0.1975 | 0.9695 | 1.77 | 2.88 |

### Top200 highly expressed (lung)

| Index | Name | P-value | Adjusted p-value | Odds Ratio | Combined score |
|---|---|---|---|---|---|
| 1 | mast cells IgE | 0.00006578 | 0.005460 | 3.37 | 32.45 |
| 2 | NK cells | 0.0002380 | 0.009875 | 2.99 | 24.97 |
| 3 | follicular B-cells | 0.01602 | 0.4408 | 2.21 | 9.12 |
| 4 | dorsal striatum | 0.02326 | 0.4408 | 2.27 | 8.54 |
| 5 | thymocyte DP CD4+CD8+ | 0.02656 | 0.4408 | 2.34 | 8.50 |
| 6 | lung | 0.03467 | 0.4796 | 2.37 | 7.96 |
| 7 | B-cells marginal zone | 0.04765 | 0.5650 | 2.20 | 6.69 |
| 8 | mast cells | 0.07935 | 0.7438 | 1.85 | 4.68 |
| 9 | mast cells IgE+antigen 1hr | 0.08066 | 0.7438 | 2.24 | 5.64 |
| 10 | adipose brown | 0.1016 | 0.8010 | 1.82 | 4.16 |

### Top200 highly expressed (muscle)

| Index | Name | P-value | Adjusted p-value | Odds Ratio | Combined score |
|---|---|---|---|---|---|
| 1 | skeletal muscle | 6.414e-10 | 5.132e-8 | 4.88 | 103.36 |
| 2 | mammary gland non-lactating | 0.000001639 | 0.00006558 | 6.93 | 92.34 |
| 3 | NK cells | 0.002315 | 0.06174 | 2.54 | 15.44 |
| 4 | heart | 0.01156 | 0.2311 | 2.32 | 10.35 |
| 5 | mast cells IgE | 0.01638 | 0.2620 | 2.20 | 9.04 |
| 6 | dorsal striatum | 0.02483 | 0.3310 | 2.24 | 8.29 |
| 7 | follicular B-cells | 0.03976 | 0.4544 | 1.97 | 6.34 |
| 8 | mast cells | 0.08351 | 0.8351 | 1.83 | 4.53 |
| 9 | macrophage bone marrow 6hr LPS | 0.1082 | 0.8763 | 1.61 | 3.58 |
| 10 | umbilical cord | 0.1101 | 0.8763 | 2.25 | 4.96 |

### Top200 highly expressed (pancreas)

| Index | Name | P-value | Adjusted p-value | Odds Ratio | Combined score |
|---|---|---|---|---|---|
| 1 | mast cells IgE | 0.001572 | 0.1336 | 2.78 | 17.93 |
| 2 | follicular B-cells | 0.004805 | 0.2042 | 2.52 | 13.46 |
| 3 | umbilical cord | 0.008128 | 0.2303 | 3.62 | 17.43 |
| 4 | mast cells | 0.01144 | 0.2431 | 2.44 | 10.90 |
| 5 | nih 3T3 | 0.06056 | 0.7649 | 2.45 | 6.86 |
| 6 | T-cells foxP3+ | 0.06749 | 0.7649 | 3.31 | 8.92 |
| 7 | cerebral cortex | 0.07294 | 0.7649 | 2.12 | 5.54 |
| 8 | Baf3 | 0.07645 | 0.7649 | 1.96 | 5.04 |
| 9 | bladder | 0.08099 | 0.7649 | 2.52 | 6.34 |
| 10 | retinal pigment epithelium | 0.1021 | 0.7690 | 1.82 | 4.15 |

### Top200 highly expressed (spleen)

| Index | Name | P-value | Adjusted p-value | Odds Ratio | Combined score |
|---|---|---|---|---|---|
| 1 | follicular B-cells | 3.490e-7 | 0.00001654 | 4.26 | 63.32 |
| 2 | NK cells | 4.297e-7 | 0.00001654 | 4.04 | 59.16 |
| 3 | mast cells IgE | 0.0002557 | 0.006563 | 3.09 | 25.59 |
| 4 | thymocyte DP CD4+CD8+ | 0.01025 | 0.1973 | 2.63 | 12.03 |
| 5 | B-cells marginal zone | 0.01895 | 0.2919 | 2.51 | 9.93 |
| 6 | bone marrow | 0.02925 | 0.3633 | 2.30 | 8.11 |
| 7 | thymocyte SP CD4+ | 0.03324 | 0.3633 | 2.92 | 9.92 |
| 8 | mast cells | 0.03775 | 0.3633 | 2.07 | 6.78 |
| 9 | nih 3T3 | 0.07094 | 0.6070 | 2.33 | 6.17 |
| 10 | T-cells CD4+ | 0.08714 | 0.6619 | 2.46 | 5.99 |

### Top200 highly expressed (testis)

| Index | Name | P-value | Adjusted p-value | Odds Ratio | Combined score |
|---|---|---|---|---|---|
| 1 | NK cells | 0.002849 | 0.1189 | 2.48 | 14.55 |
| 2 | mast cells IgE | 0.002900 | 0.1189 | 2.58 | 15.05 |
| 3 | skeletal muscle | 0.004813 | 0.1316 | 2.33 | 12.44 |
| 4 | cerebral cortex | 0.01353 | 0.2773 | 2.67 | 11.50 |
| 5 | thymocyte DP CD4+CD8+ | 0.03163 | 0.5187 | 2.26 | 7.81 |
| 6 | B-cells marginal zone | 0.05531 | 0.7533 | 2.12 | 6.14 |
| 7 | nih 3T3 | 0.07648 | 0.7533 | 2.28 | 5.85 |
| 8 | bone marrow | 0.07846 | 0.7533 | 1.95 | 4.95 |
| 9 | mast cells IgE+antigen 1hr | 0.09011 | 0.7533 | 2.16 | 5.20 |
| 10 | mast cells | 0.09219 | 0.7533 | 1.78 | 4.25 |

**Supplementary Figure 4 - Caetano et al**
