## Supplementary Data 1 for "A novel class of tiny box C/D-like RNAs reveals hidden complexity in the mouse box C/D RNA repertoire"

>tiSNORD#9778 (Gm334 intron)
gggtagcctggccttgcactcttttgttttgcatacttgg**TGACTGTGATGAATCTTACT**
**CTGAGTCA**ccagcgtatggtatgacatttgcttggtaagatatgtggaacatctgccaca
>tiSNORD#147117 (Asap1 intron)
tggtgagaagcttaaggcctccaaaggccagcagtgccttcagaacgaagccaaggtgct
ctttgcacatttcgcttcctgtcactggagtttcccaagg**GCTGTGTGATGAGTGATGGT**
**GTCTTCTCTGACAGCC**tgttaaatgtgttttggacttggaacgttttctgttttcatttc
tgttacacaaaatatcagtagatgtggctccctt
>tiSNORD#15995 (Nasr1 intron)
acacatcgcccttgctgtcttgtaaattca**GTTTATGATGAGTAGCATAATTCTGAAC**tg
aagttttaaagtactcagtgtgattgagtgaaacggtcctcagctataga
>tiSNORD#13826 (Msh3 intron)
atactttgtattggagttcttggaacaccc**TGTCTGTGATGTATAGCTGAGA**atgggtgt
cccgccacccccctcccccctctgctctgtggagttcccccatgaccctggtaggggtctc
tggagctggg
>tiSNORD#6177 (Pkig intron)
gaacagttactgcgtgtgccaagtggagcaagagaagtgtgtggca**GGGCTATGATGTGC**
**TAAGTCAGTGGATCCAGCAGAGACTCTGAGT**tcggccgctctcttagcacgtgcttgctc
cctccagttcctgtacttac
>tiSNORD#2521 (Arid4b intron)
gtcattagctggaaactaggtgttgagacacatgagcctgcgggacat**TTCACATGATGA**
**CCGTGGTCTCTGCTCAGCTGTGCTGATG**aaatgtccagttaaaatgtttgagtcatttgt
tttcaaggat
